# Assessing data quality in citizen science (preprint)

**DOI:** 10.1101/074104

**Authors:** Margaret Kosmala, Andrea Wiggins, Alexandra Swanson, Brooke Simmons

## Abstract

Ecological and environmental citizen science projects have enormous potential to advance science, influence policy, and guide resource management by producing datasets that are otherwise infeasible to generate. This potential can only be realized, though, if the datasets are of high quality. While scientists are often skeptical of the ability of unpaid volunteers to produce accurate datasets, a growing body of publications clearly shows that diverse types of citizen science projects can produce data with accuracy equal to or surpassing that of professionals. Successful projects rely on a suite of methods to boost data accuracy and account for bias, including iterative project development, volunteer training and testing, expert validation, replication across volunteers, and statistical modeling of systematic error. Each citizen science dataset should therefore be judged individually, according to project design and application, rather than assumed to be substandard simply because volunteers generated it.

## In a nutshell

- Datasets produced by volunteer citizen scientists can have reliably high quality, on par with those produced by professionals.
- Individual volunteer accuracy varies, depending on task difficulty and volunteer experience. Multiple methods exist for boosting accuracy to required levels for a given project.
- Most types of bias found in citizen science datasets are also found in professionally produced datasets and can be accommodated using existing statistical tools.
- Reviewers of citizen science projects should look for iterated project design, standardization and appropriateness of volunteer protocols and data analyses, capture of metadata, and accuracy assessment.

## Introduction

Citizen science – research that engages non-professionals in the process of creating new scientific knowledge (Bonney *et al.* 2014) – has expanded greatly in the past decade (McKinley *et al.* 2015; Figure 1). The rising interest has been fueled in part by rapid technological developments (Newman *et al.* 2012), by policy and management needs for large-scale and long-term monitoring datasets (Conrad and Hilchey 2011), and by increased emphasis on science outreach and education (Silvertown 2009). While citizen science projects vary widely in their subject matter, objectives, activities, and scale (Figures 2-4; Wiggins and Crowston 2015), one goal they share is the production of data that can be used for scientific purposes.

**Figure 1.**
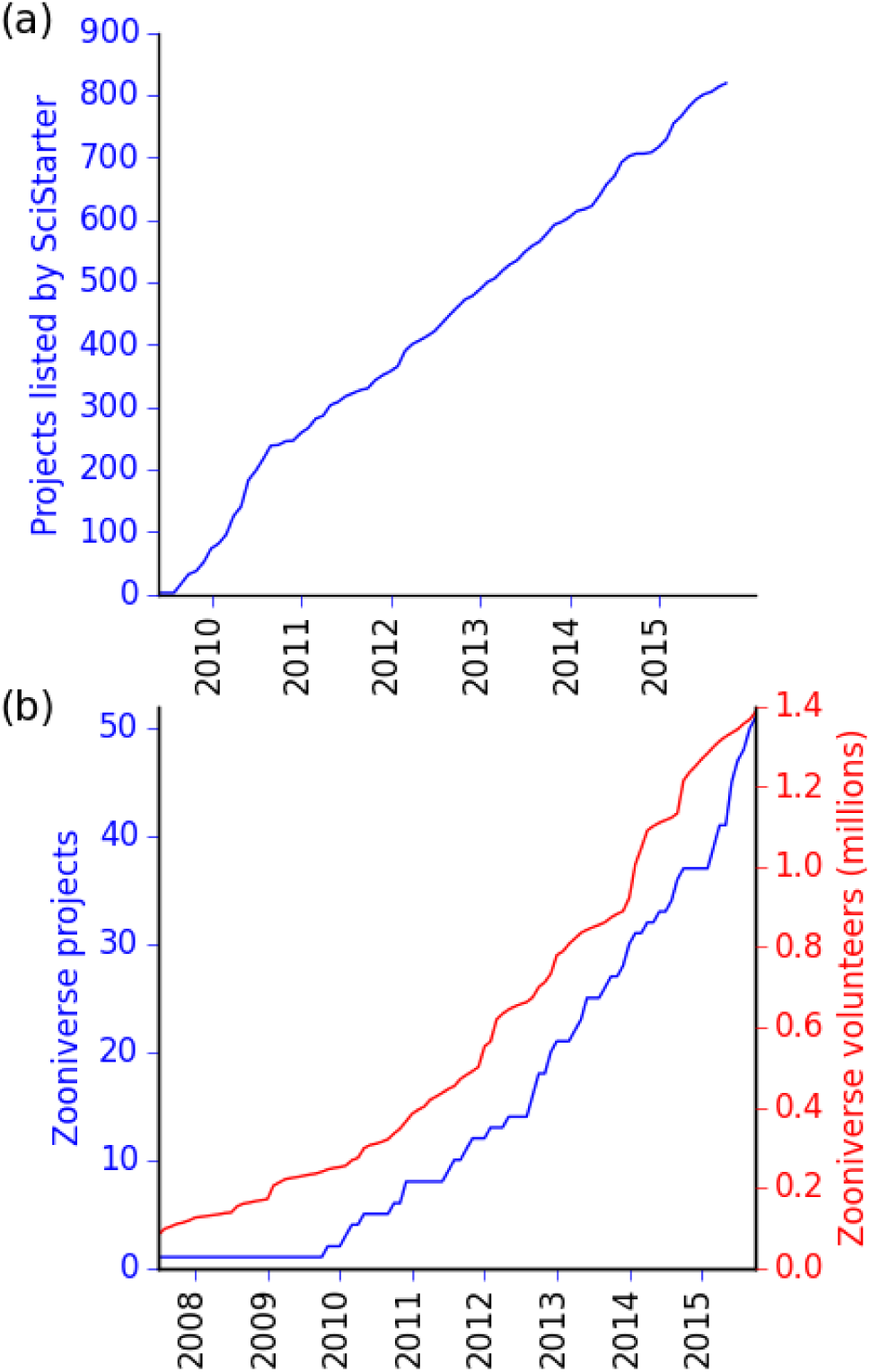
The past decade has seen a rapid increase in citizen science projects and volunteers. (a) Number of projects listed on the citizen science project directory website SciStarter, (b) Number of projects created by the citizen science portal Zooniverse (blue) and number of Zooniverse registered volunteers (red).

The ecological and environmental sciences have been leaders in citizen science, boasting some of the longest-running projects that have contributed meaningful data to science and conservation, including the Cooperative Weather Service (1890), the National Audubon Society’s Christmas Bird Count (1900, 200+ publications), the North American Breeding Bird Survey (1966, 670+ publications), the leafing and flowering times of U.S. lilacs and honeysuckles (1956, 50+ publications; Rosemartin *et al.* 2015), and the Butterfly Monitoring Scheme (1976, 100+ publications). These and other successful citizen science projects have increased ecological and environmental knowledge at large geographic scales and at high temporal resolution (McKinley *et al.* 2015). Specific advances include increased understanding of species range shifts, phenology, macroecological diversity and community composition, life-history evolution, infectious disease systems, and invasive species dynamics (Dickinson *et al.* 2010; Bonney *et al.* 2014).

Despite the wealth of information generated and the many resulting scientific discoveries, citizen science begets skepticism among professional scientists. The root of this skepticism may be that citizen science is still not considered a mainstream approach to science (Riesch and Potter 2014; Theobald *et al.* 2015). Alternatively, some professionals may believe that unpaid volunteers (hereafter, simply ‘volunteers’) are not committed or skilled enough to perform at the level of paid staff. Professional scientists have questioned the ethics of partnering with volunteers (Resnik *et al.* 2015), the “motives and ambitions” of volunteers themselves (Show 2015), and the ability of volunteers to provide quality data (Alabri and Hunter 2010). At the root of these concerns is the worry that science and policy might be derived from unreliable data. The quality of data produced by volunteers is a long-recognized concern in citizen science (Cohn 2008; Silvertown 2009; Dickinson *et al.* 2010, 2012; Conrad and Hilchey 2011; Wiggins *et al.* 2013; Bonney *et al.* 2014).

Because citizen science as a whole is often perceived as questionable science, even projects with high-quality data can find it difficult to publish results, often being relegated to educational or outreach portions of journals and conferences (Bonney *et al.* 2014). Many published peer-reviewed papers obscure the fact that citizen science data are being used by mentioning a project or database by name and citation only or by consigning the methods to supplementary materials (Cooper *et al.* 2014). Further, some parties believe that citizen science is worth more for its educational potential than for the science it can produce (Cohn 2008; Wiggins 2012). These views have made it difficult for scientists to obtain funding for potentially transformative citizen science projects (Wiggins 2012). Additionally, like many long-term projects, project leaders have often found it easier to obtain “experimental” startup funding than ongoing operational support (Wiggins and Crowston 2015).

We examine data quality practices across a wide range of ecological and environmental citizen science projects and describe the most effective methods used to acquire high-quality data. We note current challenges and future directions in assuring high-quality data. Our hope is that citizen science projects will be judged on their methods and data stewardship as a whole and not simply on whether volunteers participate in the process (Panel 1).

## What constitutes high quality data?

The concept of data quality is multi-dimensional, consisting of more than a dozen possible non-exclusive metrics (Pipino *et al.* 2002). Some metrics are task-dependent, such as timeliness of data for a particular question or objective. Other measures focus on data management practices, such as provision of relevant metadata. We focus on two objective task-independent measures of data quality that prompt the most skepticism among professional ecologists and environmental managers: accuracy and bias (Panel 1). Accuracy is the degree to which data are correct overall, while bias is systematic error in a dataset.

### Quality of data produced by professionals

A reasonable definition of *high-quality data* for citizen science is data of comparable accuracy and bias to that produced by professionals and their trainees (Bonney *et al.* 2014; Cooper *et al.* 2014; Theobald *et al.* 2015). But few projects evaluate the accuracy and bias of professionally produced data within the same contexts as volunteer-produced data. Further, much ecological data has a degree of subjective interpretation such that observations of the same sample or site vary when performed by multiple professionals or the same professional at different times.

Comparisons of data between two or more professionals can show substantial variation. For example, percentage cover estimates of intertidal communities made in 0.25-meter quadrats showed just 77.3% to 86.6% similarity (Bray-Curtis measure) between professionals (Cox *et al.* 2012). In Sweden’s National Survey of Forest Soils and Vegetation, observer identity explained nearly 20% of variance in vegetation percentage cover estimates in 100-m^2^ plots (Bergstedt *et al.* 2009). The Australian Institute of Marine Science Long-Term Monitoring Program considers newly trained professionals to be proficient once their classifications of coral reef organisms (Figure 2a) reach 90% agreement with those of established professionals (Ninio *et al.* 2003). In wildlife population surveys, it has long been acknowledged that multiple observers increase transect survey quality because of imperfect detection by single observers (Cook and Jacobson 1979). For example, under ideal conditions, single experienced observers in Alaska recorded only 68% of known moose present in aerial surveys (LeResche and Rausch 1974).

**Figure 2.**
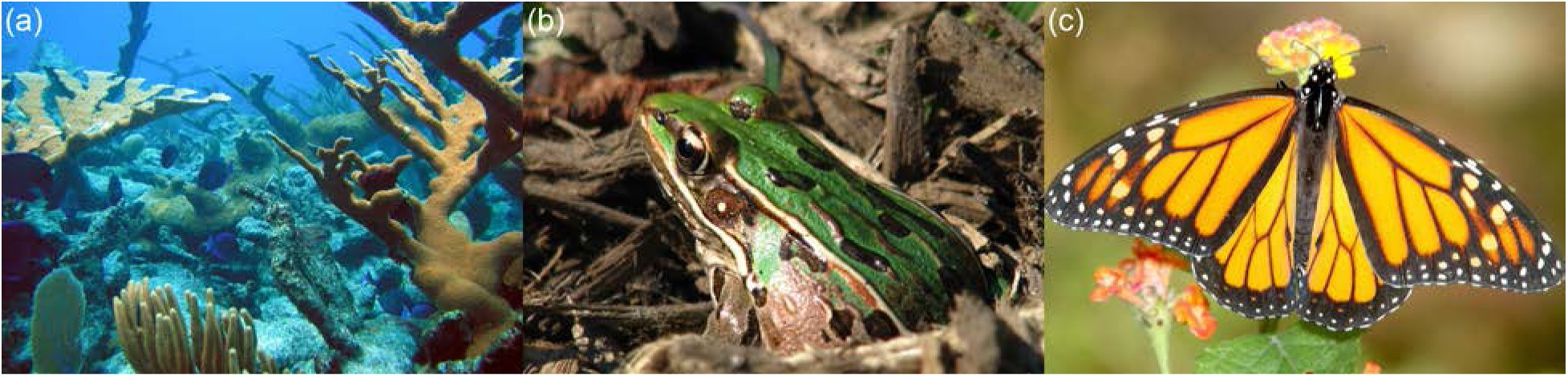
Citizen science data are collected on diverse organisms, including (a) coral reefs, such as this one in the US Virgin Islands (image by NOAA’s NOS/Flickr/public domain), (b) amphibians, such as this southern leopard frog (*Rana sphenocephala*, image by Joe McKenna/Flickr/CC BY-NC), and (c) insects, such as this monarch butterfly (*Danaus plexippus*, image by Alan Schmierer/Flickr/public domain).

Even for observations where the correct answer is more concrete, professionals sometimes make mistakes. Professionals examining trees in urban Massachusetts agreed on species identifications 98% of the time and on tree condition 89% of the time (Bloniarz and Ryan 1996). In one study recording target plant species, professionals had an 88% accuracy rate (Crall *et al.* 2011). Experts identifying large African animal species from images in Snapshot Serengeti were found to have an accuracy of 96.6%, with errors due largely to identification fatigue and data entry error (Swanson *et al.* 2016).

Because data produced by professionals and other experts can contain error and bias, comparisons between volunteer and professional data must be careful to distinguish between inter-observer variability and variability due to status as a professional or volunteer. We should also not expect the accuracy of individual volunteers to be higher than that of individual professionals.

### Quality of data produced by volunteers

Despite differences in background and experience from professional ecologists, volunteers can perform at the level of professionals for particular data gathering and processing tasks, with variation depending on task difficulty and on volunteer experience. Rates of 70-95% accuracy are typical for species identification across a diverse array of systems and taxa (Delaney *et al.* 2008; Crall *et al.* 2011; Gardiner *et al.* 2012; Fuccillo *et al.* 2015; Swanson *et al.* 2016).

Volunteers’ accuracy varies with task difficulty (Table 1). For Snapshot Serengeti, volunteers were better at identifying iconic mammals such as giraffe and zebra than less familiar mammals such as aardwolf and a set of easily confused antelope (Swanson *et al.* 2016). In anuran call surveys (Figure 2b), volunteers’ accuracy varied widely with species (Weir *et al.* 2005). The Monarch Larva Monitoring Project (Figure 2c) found reliable identification of 5th instar larvae, but not 1st and 2nd instar larvae (Prysby and Oberhauser 2004). In identifications of plant species, volunteers had an 82% accuracy rate for identification of “easy” species, but just a 65% accuracy rate for “hard” ones (Crall *et al.* 2011). Volunteers could more reliably identify street trees (Figure 3a) to genus (94% accuracy) than species (79%) (Bloniarz and Ryan 1996). Determining a crab’s species (Figure 3b) was easier (95% accuracy for seventh graders) than its sex (80% accuracy for seventh graders) (Delaney *et al.* 2008). Kelling *et al.* (2015) identified differences in bird detection (Figure 3c) and identification rates by volunteers for species that are secretive, hard to distinguish visually, or best identified by sound.

**Figure 3.**
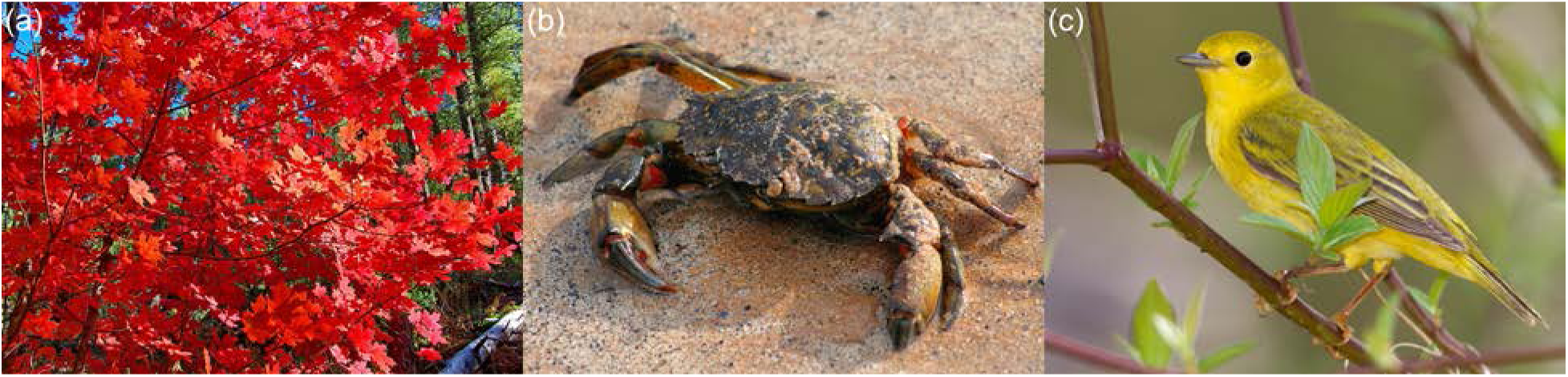
Citizen science data are collected on diverse organisms, including (a) trees, such as this red maple (*Acer rubrum*, image by Al_HikesAZ/Flickr/CC BY-NC), (b) crustaceans, such as this shore crab (*Carcinus maenas*, image by John Haslam/Flickr/CC BY), and (c) birds, such as this American yellow warbler (*Setophaga petechia*, image by Laura Gooch /Flickr/CC BY-NC-SA).

**Table 1.** Ecology and environmental citizen science task types

| Task Type | Description | Skill or training required | Examples |
| --- | --- | --- | --- |
| Taxonomic classifications | Taxonomic identification or sorting of organisms | Low to High* | **Low:** Target crab species identification (Delaney *et al.* 2008)<br>**Medium:** Antelope differentiation in Snapshot Serengeti (Swanson *et al.* 2016)<br>**High:** Cryptic bird species differentiation in eBird (Kelling *et al.* 2015) |
| Percentage cover estimates | Visual assessment of the composition of sessile organisms and/or substrate in a given area | Medium | Intertidal communities (Cox *et al.* 2012)<br>Forest vegetation (Bergstedt *et al.* 2009) |
| Presence-absence determinations | Binary determination of whether particular organisms are in a given area | Low | California Department of Fish and Wildlife's Invasive Species Citizen Science Program (www.wildlife.ca.gov/Conservation/Invasives) |
| Counts | Count of the number of individuals in a given area | Low to Medium | **Low:** Number of birds arriving at a feeder in Project FeederWatch (Bonter and Cooper 2012)<br>**Medium:** Estimating number of birds in large flocks in eBird (Kelling *et al.* 2015) |
| Organism trait measurements | Measurements of one or more traits of replicate individuals | Low to Medium** | **Low:** Plant fruiting in Nature's Notebook (Fuccillo *et al.* 2015)<br>**Medium:** Larval instar in the Monarch Larva Monitoring Project (Prysby and Oberhauser 2004) |
| Environmental measurements | Measurements of abiotic environmental conditions at a given location | Low | Atmospheric aerosols in iSPEX-EU (ispex-eu.org)<br>Precipitation in CoCoRaHS (Moon *et al.* 2009) |

\* Depending on the level of differentiation required, the familiarity of organisms, the obviousness of identifying features, and the time allowed for identification or sorting.
\*\* Depending on the trait and the instrument (if any) used to measure the trait

Volunteers often improve in accuracy as they gain experience with a project. New Snapshot Serengeti participants had an average of 78.5% accuracy, but most individuals who had classified hundreds of images had accuracies over 90% (Swanson *et al.* 2016). In the French Breeding Bird Survey, observers counted 4.3% more birds per hour after their first year of observation (Jiguet 2009), and an analysis of the North American Breeding Bird Survey also found a first-year effect (Kendall *et al.* 1996). Models of volunteers’ performance using species accumulation curves showed improvements in bird species detection and identification abilities through cumulative experience (Kelling *et al.* 2015).

## Techniques for producing high-quality ecological citizen science data

Effective methods to acquire high-quality citizen science data vary based on the type of data being created and the resources available to the project. In general, they are congruent with the procedures used by professionals (Panel 2; Wiggins and Crowston 2015). The following techniques are used by existing projects to increase the quality of citizen science data. Successful projects typically use multiple techniques.

### Iterative development of task and tool design

Iterative refinement of tasks and tools for volunteers is often a critical step in project development (Crall *et al.* 2010). The Great Sunflower Project progressively reduced the duration of observations of pollinator service, and expanded the range of plant target species, making the tasks more accessible without compromising data quality (Wiggins 2013). Mountain Watch saw a reduction in errors for hikers’ observations of alpine plant phenology (Figure 4a) when tasks and data sheets were changed to specify plots where the species were known to be present rather than any volunteer-selected location along a trail (Wiggins 2013). The Virginia Save-Our-Streams program shifted from a presence-only protocol to a count-based protocol when analyses showed that the original protocol resulted in poor data quality that consistently overrated stream condition (Engel and Voshell 2002).

**Figure 4.**
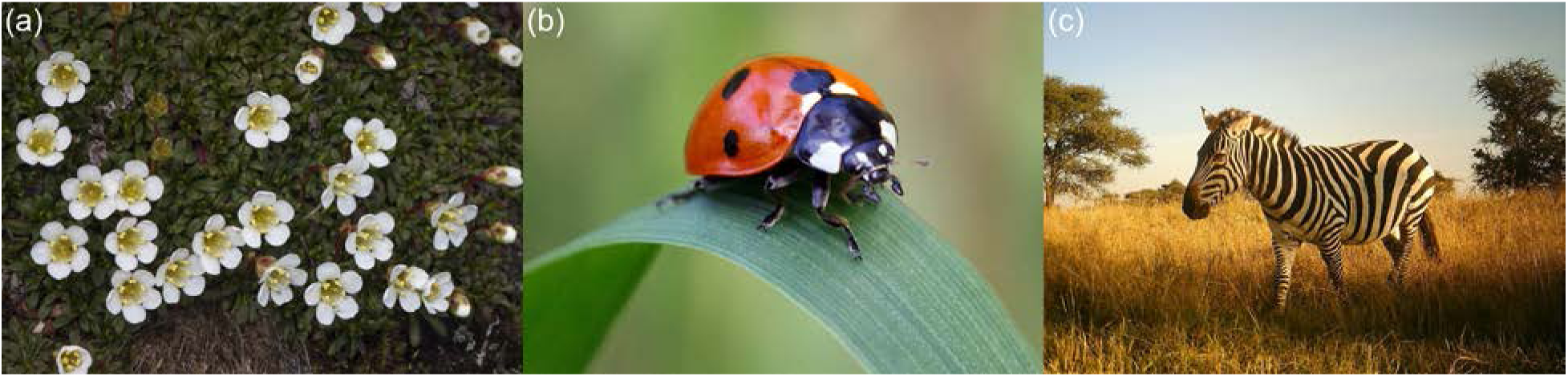
Citizen science data are collected on diverse organisms, including (a) flowering plants, such as this pincushion plant (*Diapensia lapponica*, image by Kent McFarland/Flickr/CC BY-NC), (b) insects, such as this Coccinellid (image by Stig Nygaard/Flickr/CC BY), and (c) mammals, such as this plains zebra (*Equus quagga*, image by Snapshot Serengeti).

### Volunteer training and testing

Perhaps the most obvious approach to improving data quality is to train volunteers or to require prequalification via a skills test. The Monarch Larva Monitoring Project provides volunteers an intensive training program of 4- to 11-hour workshops and focuses on long-term engagement of volunteers. Field observations and analysis of volunteer data suggest that trained and engaged volunteers produce data of similar or higher quality than hired field assistants (Prysby and Oberhauser 2004). Similarly, local volunteers monitoring tropical resources who received training over 2-3 days with shorter annual refresher training produced data of similar quality to that of professional scientists (Danielsen *et al.* 2014). Training may sometimes be self-initiated by volunteers. The Breeding Bird Survey, for example, relies upon skilled birders, who have gained their expertise over a lifetime of bird watching (Sauer *et al.* 2013).

Ongoing training can be beneficial. BeeWatch volunteers are provided ongoing feedback on their bee species identifications based on professional validation of their photographs, and this feedback increases both volunteer accuracy and retention (van der Wal *et al.* 2016). Just-in-time training can sometimes be undertaken in conjunction with project tasks. Snapshot Serengeti provides initially untrained volunteers a set of guiding filters that allows them to learn likely species identifications based on a target animal’s morphological traits (Swanson *et al.* 2016). Similarly, eBird assists its volunteers with dynamically-generated data entry forms that list the most common birds for a volunteer’s given location and time, increasing both volunteer awareness of the local species and data quality (Sullivan *et al.* 2014). Stardust@home uses known ‘seeded’ images for ongoing accuracy assessment and provides feedback to volunteers on their success rate so that they can voluntarily work on improving (Westphal *et al.* 2006).

### Use of standardized and calibrated equipment

Standardization of measurement tools and collection of instrument calibration data are common strategies for promoting high-quality data and typically mirror established professional techniques. The CoCoRaHS precipitation monitoring network requires a standardized and reliable rain gauge (Moon *et al.* 2009). Many water quality projects use standardized Secchi tubes or loan out calibrated equipment to volunteers for data capture (Sheppard and Terveen 2011), depending on the nature of the data being collected. Projects using mobile phone sensors record system data such as device model and operating system to calibrate data across devices (e.g. MyShake).

### Expert validation

When volunteers are not highly skilled or the events they observe are ephemeral, one solution is to collect vouchers allowing for expert verification. Vouchers can be physical specimens (e.g. Delaney *et al.* 2008; Gardiner *et al.* 2012) or photographs, video, or audio recordings (Kageyama *et al.* 2007). The eMammal project asks volunteers to set up motion-triggered cameras to monitor North American mammals (McShea *et al.* 2015). These volunteers make species identifications for “their” images, while the images themselves serve as vouchers, allowing experts to validate species identifications. Expert validation of volunteer classifications has been shown to be more cost-effective than direct expert classification for lady beetles (Figure 4b; Gardiner *et al.* 2012).

However, expert validation of every data point can be impractical, and for large projects, efficiently targeting likely wrong answers is key. Project FeederWatch uses a “smart filter” system that flags observations of unlikely species and unusually large numbers of birds. Flagged data are immediately sent to regional experts who then ask for photographic vouchers and supporting details from volunteers to validate the sighting. Over three years, just 1.3% of observations required expert review (Bonter and Cooper 2012). Similarly, Snapshot Serengeti uses a suite of post-hoc statistical metrics to identify “difficult” images of African animals to be sent for expert review (Swanson *et al.* 2016).

### Replication and calibration across volunteers

Some projects make multiple independent measurements for each subject to improve data quality. Projects on the Zooniverse platform show each digital voucher to multiple volunteers, with all resulting classifications combined into a “consensus” answer. For example, each image in Snapshot Serengeti (Figure 4c) is shown to 5-25 volunteers and its consensus answer is the plurality of identifications from all volunteers. Consensus improved accuracy from 88.6% to 97.9% over single classifications (Swanson *et al.* 2016).

When replication for all data points is not practical, calibration across volunteers using targeted replication allows for statistical control of data quality. In Mountain Watch, volunteers collect data at fixed locations as well as self-selected locations, with trained staff also reporting data from the fixed plots; this permits verification of observations from volunteers against those of staff naturalists. The fixed plots also allow for statistical normalization across volunteers and additional logger data from these plots provide covariates for data analysis (Wiggins 2013). Another calibration technique involves injecting professionally-classified (e.g. Stardust@home; Westphal *et al.* 2006) or artificially generated (e.g. Planet Hunters; Schwamb *et al.* 2012) vouchers into voucher sets given to volunteers for classification in order to evaluate ongoing volunteer performance.

### Skill-based statistical weighting of volunteer classifications

Methods are emerging for weighting volunteer classifications based on individual characteristics, such as skill level. For projects with multiple classifications per captured datum, volunteer skill can be assessed via frequency of agreement with other volunteers. For Snapshot Serengeti data, weighting increased consensus accuracy from 96.4% to 98.6% (Hines *et al.* 2015). In cases where there is only one classification per captured datum, skill can be assessed by testing or other means. The observation skill of eBird users was assessed using species accumulation curves, and when skill was incorporated into bird species distribution models, model accuracy increased for approximately 90% of the 120 species tested (Kelling *et al.* 2015).

### Accounting for random error and systematic bias

Data produced through citizen science may contain error and bias, but existing statistical and modeling tools can accommodate these errors and biases to produce meaningful inference. A common concern is that citizen science data is too “noisy”: it has too much variability. For some projects, collecting a sufficiently large amount of data may be enough to reduce non-systematic error in volunteer-produced data through the law of large numbers (Bird *et al.* 2014). eBird data accumulate at the rate of millions of observations monthly (Sullivan *et al.* 2014), and the resulting range maps and temporal distribution patterns concur with professional knowledge (Wiggins 2012). Similarly, with more than 750,000 individual reports, the U.S. Geological Survey’s “Did You Feel It?” program yields highly accurate measures of earthquake strength when compared with ground sensors (Atkinson and Wald 2007).

Many of the systematic biases in citizen science data are the same biases that occur in professionally collected data: spatially and temporally non-random observations (biased by things such as time of day or week, weather, and human population density; e.g. Courter *et al.* 2013), non-standardized capture or search effort, under-detection of organisms (Elkinton *et al.* 2009; Crall *et al.* 2011), confusion between similar-looking species, and the over- or under-reporting of rare, cryptic, or elusive species compared to more common ones (Gardiner *et al.* 2012; Kelling *et al.* 2015; Swanson *et al.* 2016). However, all these biases are also found in professional ecological research, and there are many methods for statistically controlling for and modeling these biases, as long as the relevant metadata are recorded (Bird *et al.* 2014).

The only known bias specific to citizen science is the potentially high variability among volunteers in terms of demographics, ability, effort, and commitment. Modeling characteristics that vary among volunteers such as age, previous experience, formal education, attitudes, and training methods may increase data reliability, although the magnitude of the effect may be project- or task-dependent (Galloway *et al.* 2006; Delaney *et al.* 2008; Crall *et al.* 2011). Bird *et al.* (2014) thoroughly describe existing statistical methods – such as generalized linear models, mixed-effect models, hierarchical models, and machine learning algorithms – that can be used to properly analyze large and variable datasets produced by citizen science projects.

## Challenges and the future of high-quality citizen science data

Technology is rapidly developing to make implementing best practices for high-quality citizen science data easier, but challenges in project technologies and data management still remain. Online resource sites (e.g. Cornell’s Citizen Science Toolkit, U.S. Federal Crowdsourcing and Citizen Science Toolkit), platforms for building online citizen science projects (e.g. Zooniverse Project Builder, CrowdCrafting), and data entry tools for field data (e.g. iNaturalist, CitSci.org, iSpot) are making it easier than ever to build citizen science projects with online components. However, research in the field of human-computer interaction is beginning to show direct and indirect impacts of online project and technology design on volunteer performance (Bowser *et al.* 2013; Eveleigh *et al.* 2014), and more such research is needed. The next generation of multipurpose data entry platforms should allow for customized data constraints and real-time outlier detection to reduce data entry error. Additionally, repositories to support terabyte-scale multimedia voucher sets are increasingly needed (e.g. McShea *et al.* 2015). Other technological challenges include unreliable mobile device GPS performance, necessity of offline functionality for mobile devices, issues of usability and accessibility, and user privacy protections (Bowser-Livermore and Wiggins 2015; Wiggins and He 2016).

Additional research is also needed in the application of existing statistical and modeling tools to citizen science datasets, which sometimes present additional challenges (Bird *et al.* 2014). Currently, analyses of complex citizen science data often require custom solutions developed by professional statisticians and computer scientists, using high performance or cloud computing systems (e.g. Yu *et al.* 2010; Hochachka *et al.* 2012) – resources that are not available to most projects. Generalizable and scalable methods to analyze variable spatiotemporal datasets will be increasingly valuable, and borrowing techniques from other fields may prove beneficial. The information science field has developed sophisticated methods for combining categorical classifications across multiple observers (e.g. Woźniak *et al.* 2014). Similarly, the social sciences have developed reliability and aggregation metrics that can be adapted to accommodate heterogeneous volunteer data. In the computer science field, optimal crowdsourcing has commercial applications, engendering new human computation journals and conferences (e.g. Human Computation Journal, AAAI Human Computation conference). Task allocation algorithms, in particular, have the potential to improve both data quality and project efficiency by routing content to the best individuals (Kamar *et al.* 2012).

## Conclusion

As citizen science continues to grow and mature, we expect to see a growing awareness of data quality as a key metric of project success. Appropriate metrics of data quality compare data produced by volunteers against similar data produced by professionals and distinguish inter-observer variability from variability due to observer experience. Evidence from across a diverse range of different task types and study systems shows that volunteers can produce high-quality data, and that accuracy is particularly high for easy tasks and for experienced volunteers. High-quality data can be produced using a suite of techniques, and investment in additional research and technology has the potential to augment these techniques and make them more broadly accessible. We suggest that Panel 1 be used as a guide by citizen science evaluators, project creators, and data users as a standard to gauge data quality. As we face grand challenges related to global environmental change, citizen science emerges as a general tool to collect otherwise unobtainable high-quality data in support of policy and resource management, conservation monitoring, and basic science.

## Acknowledgements

MK is supported by a grant from the National Science Foundation, through the Macrosystems Biology Program (award EF-1065029). BS acknowledges support from Balliol College, Oxford, and the National Aeronautics and Space Administration (NASA) through Einstein Postdoctoral Fellowship Award Number PF5-160143 issued by the Chandra X-ray Observatory Center, which is operated by the Smithsonian Astrophysical Observatory for and on behalf of NASA under contract NAS8-03060.

## Panel 1. Questions to consider when evaluating citizen science projects for data quality

The following questions are based on existing research and are meant for use by creators, evaluators, and users of citizen science data. Creators of citizen science projects may use them to guide project development, and are encouraged to reference them in project methods. Evaluators and reviewers of citizen science proposals and manuscripts may use them to better gauge the quality of data in citizen science projects. And citizen science data consumers may use these questions to ascertain suitability of datasets for particular scientific questions. Future research should build on current knowledge to strengthen and broaden best practices for data quality.

### Does the project use iterative design?

Developing tools and protocols for a project that produces high-quality data requires iteration, using one or more rounds of pilot or beta testing to ensure a procedure volunteers can perform successfully without confusion or systematic errors.

### How easy or hard are the tasks?

Easy tasks likely have high accuracy with little bias. Hard or complex tasks may require additional effort on the part of the project managers to promote accuracy and account for bias. Such efforts include training, pre-tests, ongoing volunteer assessment, expert validation, classification replication, and application of statistical tools.

### How systematic are the task procedures and data entry?

High-quality data requires straightforward and systematic capture, classification, and data entry procedures for the volunteers to follow. For online data entry, fields should enforce type (e.g. counts must be integers) and for categorical variables, users should select from lists rather than entering free-form text.

### What equipment are volunteers using?

Any equipment used for measurements should be standardized and calibrated across volunteers.

### Does the project record relevant metadata?

Projects should record metadata that may influence volunteer data capture or collection. Such data might include environmental conditions (temperature, precipitation, time of day, etc.), equipment or device settings (such as mobile device OS version), or characteristics of the volunteers themselves (such as level of education or training). If characteristics of volunteers are collected, project managers should seek approval from the relevant human subjects review board. Projects should also retain volunteer identifiers (anonymized if necessary). These metadata can be used to statistically model bias to increase valid inference from project data.

### Is collection effort standardized or accounted for in data analysis?

Standardized effort (capturing data at specified places, times, and/or durations of time) is ideal for ensuring unbiased data. However, many projects cannot standardize effort; for these projects, it is imperative that effort is reported by volunteers and is accounted for in statistical models and analysis.

### Does the project assess data quality by appropriate comparison with professionals?

In reporting results, citizen science projects should compare volunteer data accuracy with that of professionals. Importantly, between-professionals accuracy should also be assessed so that variation due to individuals is not confused with variation due to volunteer-professional differences.

### Is the data appropriate for the project’s management objectives or research questions?

In particular, data should be of sufficient quantity and should cover timescales and geographic extents commensurate with project objectives. Data may also need to be timely, depending on the application.

### Are good data management practices used?

Citizen science project managers should implement best practices in data management (e.g. (Borer *et al.* 2009; Michener and Jones 2012; Wiggins *et al.* 2013). In particular, data should be stored securely in a consistent and concise format that is easy to interpret and use and is made accessible to data users.

## Panel 2. Data capture and data classification

We distinguish between data *capture* (collection and observation) and data *classification* (the interpretation of raw data into an analyzable form). An example of data *capture* is the collection of insects by pitfall trap. The corresponding data *classification* is the determination of their taxonomic identifications. These two steps are frequently conducted concurrently by professionals (e.g. percentage cover estimates), but separating the process into discrete tasks allows better control over statistical analysis of data error. In citizen science projects, volunteers may conduct data capture, data classification, or both.

### Volunteer capture, professional classification

Volunteers collect samples and send them to professionals for analysis. This method is typically employed to gather data at large spatial scales and when laboratory methods are required. *Examples*: Clean Air Coalition of Western New York, Lakes of Missouri Volunteer Program, American Gut

### Professional capture, volunteer classification

Professionals select subjects to evaluate, but lack capacity to classify all subjects. Projects that use large volumes of digital images produced by cameras set up by experts fall into this category. *Examples*: Snapshot Serengeti (camera traps), Floating Forests (satellite imagery), Season Spotter (automated near-earth cameras).

### Volunteer capture and classification

Volunteers collect samples, make observations, or set up automated collection devices. They also classify the observations, samples, or vouchers. *Examples*: Project Feeder Watch, eBird, eMammal, Monarch Larva Monitoring Project, Nature’s Notebook

